# Sparse coding at the input stage of the hippocampus enables fast learning

**DOI:** 10.64898/2026.04.27.721238

**Authors:** Animesh Sasan, Robert M. Mok

## Abstract

The hippocampus is crucial for fast learning, a capacity commonly attributed to the sparse code of the dentate gyrus (DG), the input stage of the trisynaptic circuit. However, this has been assumed rather than demonstrated, and we lack a principled account of when sparsity helps or hurts learning and what is traded off along the sparse-to-dense continuum. We characterize sparse coding in neural networks trained on nonlinear tasks and show that the level and position of sparsity determine learning speed and the memorization–generalization trade-off. Learning speed follows an inverted-U function, peaking at capacity-sufficient codes. Critically, sparsity must be implemented at the input stage, mirroring the DG. Sparsity speeds learning by producing high-dimensional, pattern-separated representations, but at a cost: models learn through memorization and fail on new examples. Our work establishes principles linking sparsity to learning, casting the DG’s sparse code as a mechanism for episodic snapshots rather than general structure.

## 2 Introduction

The hippocampus has long been known to be essential for fast learning and one-shot memory formation([1]; [2]; [3]; [4]). This capacity is thought to be enabled by sparse coding in the dentate gyrus (DG), which supports pattern separation by reducing overlap between similar events and thereby minimizing interference([5]; [6]; [7]; [8]; [9]). Despite an extensive body of work on the hippocampus, there is surprisingly little research characterizing the functional properties of the sparse code of the DG, and the benefits of sparse coding have been questioned more broadly([10]). A large majority of work has focused on modeling the associative aspects of memory function in CA3 using auto-associative networks([11]; [12]; [13]; [14]), CA1 as a comparator([15];[16]), and the roles of the trisynaptic and monosynaptic pathways([17]; [18]). Models that incorporated the DG mainly highlighted its ability to orthogonalize patterns for effective memory formation in downstream regions([14]), but did not systematically evaluate the computational properties of the DG sparse code nor its role in complex cognition.

An emerging line of research suggests that the hippocampus is not only involved in memory but plays a key role across a wide range of cognitive domains from perception([19]; [20]) to complex goal-oriented behavior([21]; [22]; [23]; [24]). Theoretical work has largely focused on memory (recall based on incomplete inputs;[25]; [26]) or simple linear tasks that are amenable to mathematical analysis([27]). However, there is growing recognition of the critical need for models that actually perform complex (e.g., nonlinear) tasks and capture intelligent human behavior([28]; [29]). What is the DG useful for in complex cognition? Pattern separation supports not only for memory but challenging perceptual tasks([20]). It may also be important for solving complex, non-linear tasks that require breaking down the problem into sub-components to facilitate building a cognitive map for effective behavior([21]).

Memory for specific events (e.g., seeing a specific dog) and generalized knowledge (e.g., the concept of dog) sit on two extremes of the memorization–generalization axis. The hippocampus is traditionally associated with specific memories, and the cortex — typically modeled with a deep neural network (DNN) with a dense code([30]; [31]) — is thought to be responsible for generalized, semantic knowledge about the world([32]; [33]; [34]). In machine learning, DNNs are not typically trained with explicit sparsity constraints on units, which likely contributes to their slow, sample-inefficient learning that requires thousands of examples to solve a task. Because every input drives the entire network, weight updates from different inputs interfere with one another. Notably, this is not addressed by standard sparsity regularization. L1 and related penalties act on weights rather than units, and the constraint applies uniformly across the network, so every input still engages the whole network. They also *slow* learning, since they impose an additional objective that competes with the task. Despite these costs of dense coding, model overparameterization — having more units and connections than necessary for the training task — is central to high generalization performance in deep learning([35]; [36]; [37]), which demonstrates the power of dense networks. By contrast, sparse coding, often heralded as a hallmark of effective and efficient neural computation([38]; [39]), constrains activity rather than connectivity, and may be key to fast learning.

To what degree and at what position in the processing stream does a sparse code facilitate fast learning, and is there a limit to its capacity? Does fast, sparse learning come at the expense of generalized knowledge, or can a sparse code support both? Motivated by these considerations, we used an artificial neural network approach and conducted a systematic investigation of how varying levels of hippocampal-like sparsity at different processing stages affect learning speed, generalization performance, and internal representations on complex non-linear tasks, including the 2-dimensional exclusive-or (XOR) task and a 7-dimensional (7D) category learning task([40]). The 7D task requires distinguishing inputs that overlap heavily in their features, including exceptions that share most features with the opposite category.

## 3 Results

### 3.1 Fast learning emerges at sparse, capacity-sufficient codes

The hippocampus’ capacity for fast learning is attributed to the highly sparse code of the DG. However, extreme sparsity constrains representational capacity, whereas an overly dense code requires more training to resolve interference between overlapping representations. Therefore, we hypothesized that fast learning is optimized with a relatively high level of sparsity that is nevertheless capacity-sufficient for the task.

To assess the effect of sparsity level on learning speed, we trained two- and three-layer networks on a two-dimensional XOR task while systematically varying the level of sparsity in every layer and assessed their learning performance. Sparsity was implemented using a *k*-winners-take-all mechanism that restricts activity in a layer to the top-*k* most strongly activated units. For instance, with 200 units per layer and 5% sparsity, only the top 10 most strongly activated units remain active (*k* = 10) for a given input, while the remaining are inhibited by setting their activations to zero. Sparsity denotes the proportion of units active for a given input, so lower values indicate sparser codes. All models were trained trial-by-trial with a binary cross-entropy objective and assessed using classification accuracy on all input exemplars, averaged across 50 runs (see Methods for details).

Across architectures, sparse models learned faster than the dense model (Fig.1B-C). Notably, the 40% sparse model exhibited learning dynamics comparable to the dense baseline, whereas sparser models trained substantially faster. The 5% sparse models learned fastest (number of trials to a 95% accuracy threshold). The two-layer 5% sparse model required only ∼ 8 trials on average (two stimulus repetitions) to reach 95% accuracy (Fig.1C). Deviations from this sparsity level in either direction led to slower learning. In particular, the 1% sparse model required ∼ 28 trials to reach threshold, comparable to the dense model, which required ∼ 30 trials. Our results reveal a non-monotonic relationship between sparsity and learning speed, where the sparsity level trades off representational capacity against learning efficiency. The fastest-learning models fall within the sparsity range estimated for the DG([41]; [42]). Models trained with random-*k* sparsity failed to learn the task, demonstrating that our results are not due to sparsity alone, but from effective feature selection enabled by top-k’s activation-dependent mechanism (Fig.S4).

**Figure 1:**
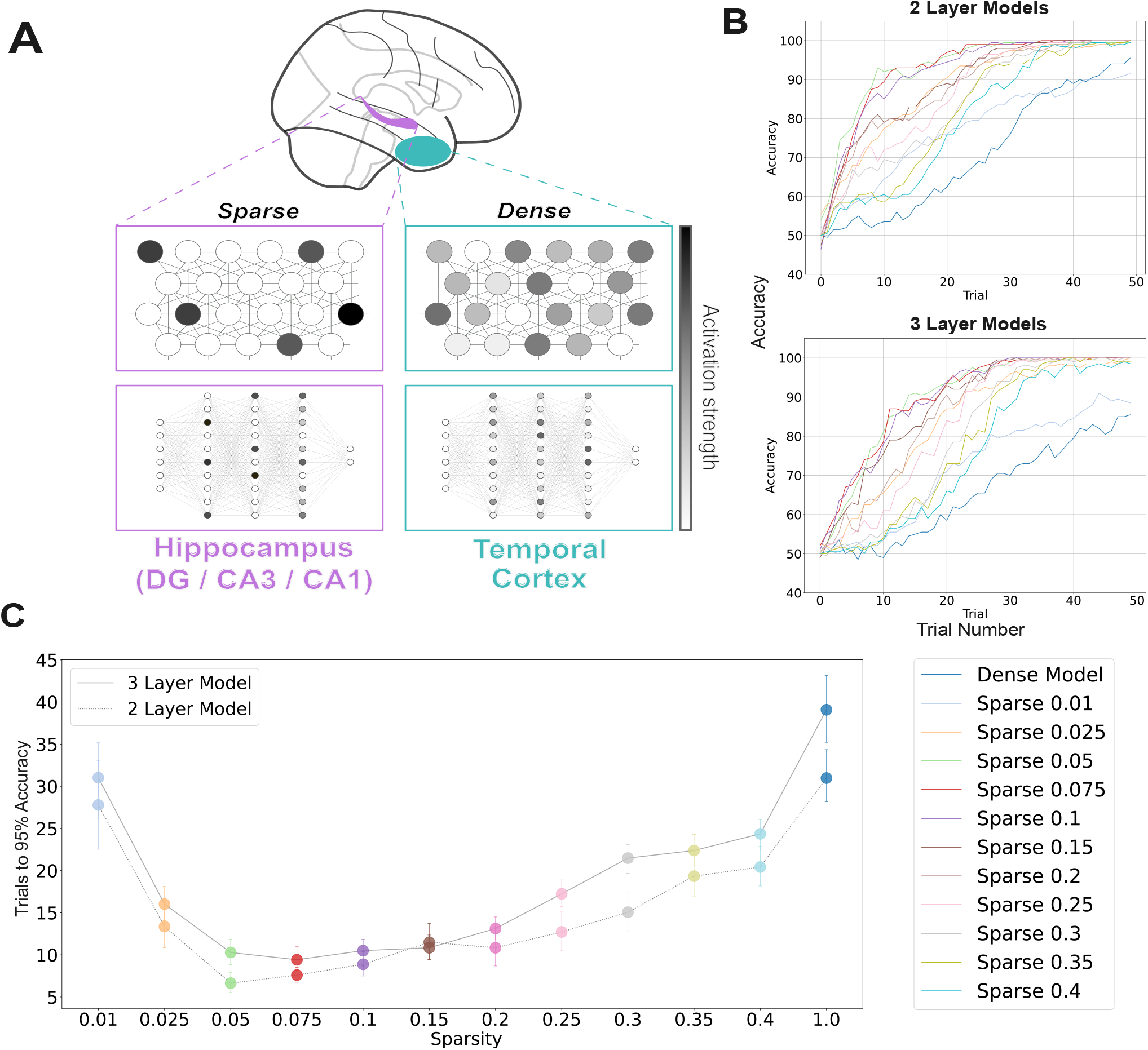
Hippocampus-like sparsity produces fast learning. (A) Illustration of hippocampal sparse representations (purple) contrasted with a distributed, dense cortical code (anterior/inferior temporal cortex; teal). Conceptual illustration of sparse and dense codes (top) and an example of their implementation in artificial neural networks (bottom). (B) Learning curves for two-layer (top) and three-layer (bottom) networks trained on an XOR task under varying sparsity levels showing fast learning for a range of sparsity levels. Sparsity denotes the proportion of neurons active within each layer for a given input. (C) Mean number of trials required to reach 95% accuracy as a function of sparsity level. Models exhibit a non-monotonic dependence on sparsity, with fastest learning at approximately 5% sparsity. Error bars represent 95% bootstrapped confidence intervals (CIs) of the mean.

One question is whether the size of the network and the number or proportion of active units matters for sparsity-based fast learning. While wider networks provide greater representational capacity, they also introduce more trainable parameters (weights), which could slow learning. Moreover, under percentage-based sparsity, increasing width increases the number of active units (*k*), raising the question of whether learning depends on *k* itself or on the sparsity ratio. To address this, we trained two- and three-layer networks with varying hidden-layer widths (100, 200, 300, 400, 1000) under two sparsity regimes: percentage-based sparsity (top 5% active units) and absolute sparsity (top-*k* with *k* = 10, corresponding to 5% for 200-unit wide networks; see Methods). We found that learning speed was relatively uniform across models of varying widths (Fig.S1). For sparse models, there was a slight reduction in learning speed for the smallest models (100 units), likely due to limited capacity (*k* = 5). Otherwise, learning speed was comparable across widths, with learning speed slightly faster for 300-1000 unit wide models, for both models with percentage (5%) or a fixed sparsity (*k* = 10). Importantly, increasing width does not systematically increase learning speed in dense architectures. In contrast, sparse networks maintain efficient learning even as model capacity grows.

### 3.2 First-layer sparsity is necessary for fast learning

Next, we investigated how the position of sparsity within the network shapes learning dynamics with sparsity fixed at 5%. In particular, we asked whether input-stage sparsity — analogous to the DG sparse code in the hippocampal circuit — is required for fast learning, or whether learning speed is insensitive to where sparsity is implemented.

We found that learning was faster in all models with input-stage (first layer) sparsity, mirroring the DG-CA3-CA1 pathway. In contrast, models in which sparsity was implemented in later layers (second- or third-layer sparse models) exhibited substantially slower learning dynamics (Fig.2(A-B)).

First-layer sparse and complete sparse models reached 95% accuracy in ∼8 trials (Fig.2(B)). In comparison, second-layer sparse models required ∼25 trials, while the fully dense model reached this threshold only after ∼30 trials. All architectures containing sparse layers trained faster than the dense baseline. These results indicate that learning speed depends not only on the presence of a sparse code but on its position within the network. As above, models trained with random-*k* sparsity restricted to specific layers also failed to learn (Fig.S4(B)).

**Figure 2:**
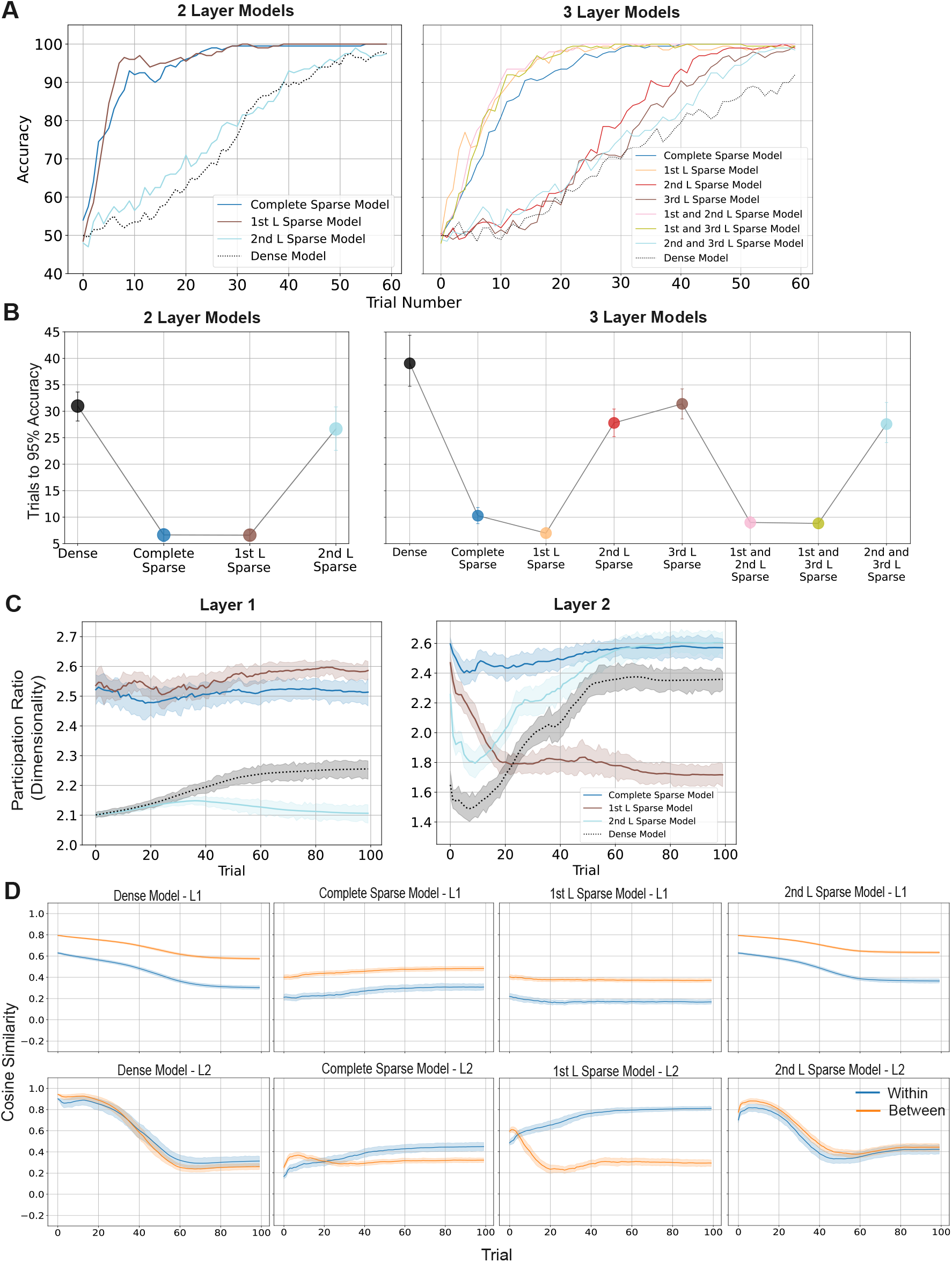
Position of sparsity determines learning speed and representational separation. (A) Learning curves for networks with sparsity implemented at different layers, showing fastest learning with first-layer sparsity. (B) Number of trials to reach 95% accuracy across models. Models with a sparse first layer learn fastest, whereas the second-layer and third-layer sparse models learn more slowly. Error bars represent 95% bootstrapped CIs. (C) Dimensionality (participation ratio; PR) learning dynamics for the first and second hidden layers for two-layer networks. (D) Representational dynamics over learning reveal rapid task-relevant separation in later layers. Cosine similarity between activations for within-class and between-class input pairs at each layer, for two-layer models. First-layer sparse and complete sparse models show greater within-class than between-class similarity in layer 2. Shaded error bars represent 95% bootstrapped CIs.

### 3.3 Sparsity speeds learning by producing high-dimensional, pattern-separated representations

Why does first-layer sparsity speed learning? We hypothesized that, as in the DG in the trisynaptic circuit, first-layer sparsity in our models (e.g., first-layer and first-and-second-layer sparse) implements pattern separation by orthogonalizing activation patterns, enabling fast learning. Since this would increase the number of distinct activation patterns across inputs, model representations should exhibit higher dimensionality. For a measure of the models’ effective dimensionality over learning, we computed the participation ratio (PR) across all input activation patterns and model layers([43]; see Methods).

In the first layer, first-layer sparse and complete sparse models exhibited markedly higher dimensionality than for second-layer sparse (which has a dense first layer) and dense models, reflecting more orthogonalized, pattern-separated representations (Fig.2C, left). This pattern remained stable over learning. In the second layer, first-layer sparse models exhibited high dimensionality which dropped sharply early in learning (Fig.2C, right). Notably, the complete sparse model dimensionality remained high, as sparsity is enforced in this layer. Second-layer sparse and dense models started with a lower dimensionality, reflecting more overlapping representations.

To probe how differences in dimensionality are reflected in the geometry of task representations, we analyzed how representational similarity across categories develops over learning. For each model and layer, we computed the cosine similarity between the unit activations for all input pairs, then plotted the mean similarity for within-class and between-class pairs. There were marked differences in the second layer across models, with only minor differences in the first layer (Fig.2(D)). In the second layer, the dense and second-layer sparse models exhibited higher initial similarity across both within- and between-category pairs, likely due to more overlapping representations early in learning. First-layer and complete sparse models exhibited greater within-class than between-class similarity early on in learning, corresponding to their learning speed. This separation was strongest for the first-layer sparse models. These results also mirror dimensionality compression reflected in the first-layer sparse models’ second layer, where dimensionality is initially high, but then quickly reduces to a lower level. In contrast, the complete sparse model’s PR remain high due to the enforced sparsity at the second layer. These results indicate that first-layer sparsity accelerates the emergence of separated class representations and leads to substantially faster learning through orthogonalized representations – at least for the examples presented during learning (i.e., the training set).

Why does the first-layer sparse model compress information more than other models while maintaining high accuracy? Does this reflect efficient learning, or simply memorization without generalization? Since the current XOR task has only four training examples and no held-out test set, we cannot distinguish between memorization and generalization. To resolve this, we turn to a higher-dimensional task in the next section.

### 3.4 Sparsity level and position jointly set the memorization–generalization trade-off

The hippocampus is traditionally associated with episodic memory, which sits closer to “memorization” than generalized knowledge. Indeed, a localist code — an extreme sparse code where only one unit is active for one input (or concept) — would store individual experiences rather than learn general knowledge about the world. If sparsity speeds learning by sacrificing generality for specificity, the DG’s sparse code should sit closer to memorization. Here, we tested whether and how sparsity determines the trade-off between memorization and generalized knowledge acquisition. To investigate this, we trained networks on a complex 7D, non-linear category learning task (with a train–test split) from the human category-learning literature, developed to study how humans learn and generalize categories with complex yet realistic category structure([40]). Following the same procedure as above, we trained neural network models to assess how sparsity relates to the trade-off between memorization and generalization. All models were trained on a trial-by-trial basis on the training set (20 exemplars), and generalization performance was evaluated on the held out test set (6 exemplars) after every training trial (see Table 1 for the full category structure).

**Table 1:** 7D category learning task. Each input consists of seven binary features (*a*–*g*) and a category label. We indicate whether each example belongs to the training or test set. This is a family resemblance category structure, and it can also be considered a rule-plus-exception structure (see original paper for details;[40])

| $a$ | $b$ | $c$ | $d$ | $e$ | $f$ | $g$ | Category | Split |
| --- | --- | --- | --- | --- | --- | --- | --- | --- |
| 1 | 1 | 0 | 0 | 1 | 1 | 0 | 1 | Train |
| 0 | 0 | 1 | 0 | 0 | 0 | 1 | 1 | Train |
| 0 | 0 | 0 | 1 | 0 | 0 | 1 | 1 | Train |
| 0 | 0 | 1 | 1 | 0 | 0 | 0 | 2 | Train |
| 1 | 1 | 1 | 0 | 1 | 1 | 1 | 2 | Train |
| 1 | 1 | 0 | 1 | 1 | 1 | 1 | 2 | Train |
| 1 | 1 | 1 | 1 | 1 | 1 | 0 | 1 | Train |
| 0 | 0 | 1 | 1 | 1 | 1 | 0 | 1 | Train |
| 1 | 1 | 0 | 1 | 0 | 1 | 0 | 1 | Train |
| 1 | 0 | 1 | 1 | 1 | 0 | 0 | 1 | Train |
| 0 | 1 | 1 | 0 | 1 | 1 | 0 | 1 | Train |
| 1 | 0 | 0 | 0 | 0 | 0 | 1 | 1 | Train |
| 0 | 1 | 0 | 0 | 0 | 0 | 1 | 1 | Train |
| 0 | 0 | 0 | 0 | 0 | 0 | 0 | 2 | Train |
| 0 | 0 | 0 | 0 | 1 | 1 | 0 | 2 | Train |
| 0 | 1 | 0 | 1 | 0 | 0 | 0 | 2 | Train |
| 1 | 0 | 0 | 0 | 1 | 0 | 0 | 2 | Train |
| 0 | 0 | 1 | 0 | 0 | 1 | 0 | 2 | Train |
| 1 | 1 | 1 | 1 | 1 | 0 | 1 | 2 | Train |
| 1 | 1 | 1 | 1 | 0 | 1 | 1 | 2 | Train |
| 0 | 1 | 1 | 1 | 0 | 1 | 0 | 1 | Test |
| 1 | 0 | 1 | 0 | 1 | 1 | 0 | 1 | Test |
| 1 | 1 | 0 | 1 | 1 | 0 | 0 | 1 | Test |
| 0 | 1 | 0 | 0 | 0 | 1 | 0 | 2 | Test |
| 0 | 0 | 1 | 0 | 1 | 0 | 0 | 2 | Test |
| 1 | 0 | 0 | 1 | 0 | 0 | 0 | 2 | Test |

Examining the effect of sparsity level on learning speed in complete sparse models, we found the same non-monotonic relationship between sparsity and learning speed on the training set, with 5-15% sparsity models learning the fastest (Fig.3 A). Interestingly, test accuracy – reflecting generalized knowledge – reveals a markedly different pattern. Test accuracy for sparse models hovered slightly above chance level (50-55%), and denser models showed increasingly better generalization performance (Fig.3 B), suggesting that sparse models learned fast by memorizing inputs rather than learning the general task structure. These results demonstrate a trade-off between memorization and generalization across sparsity levels.

**Figure 3:**
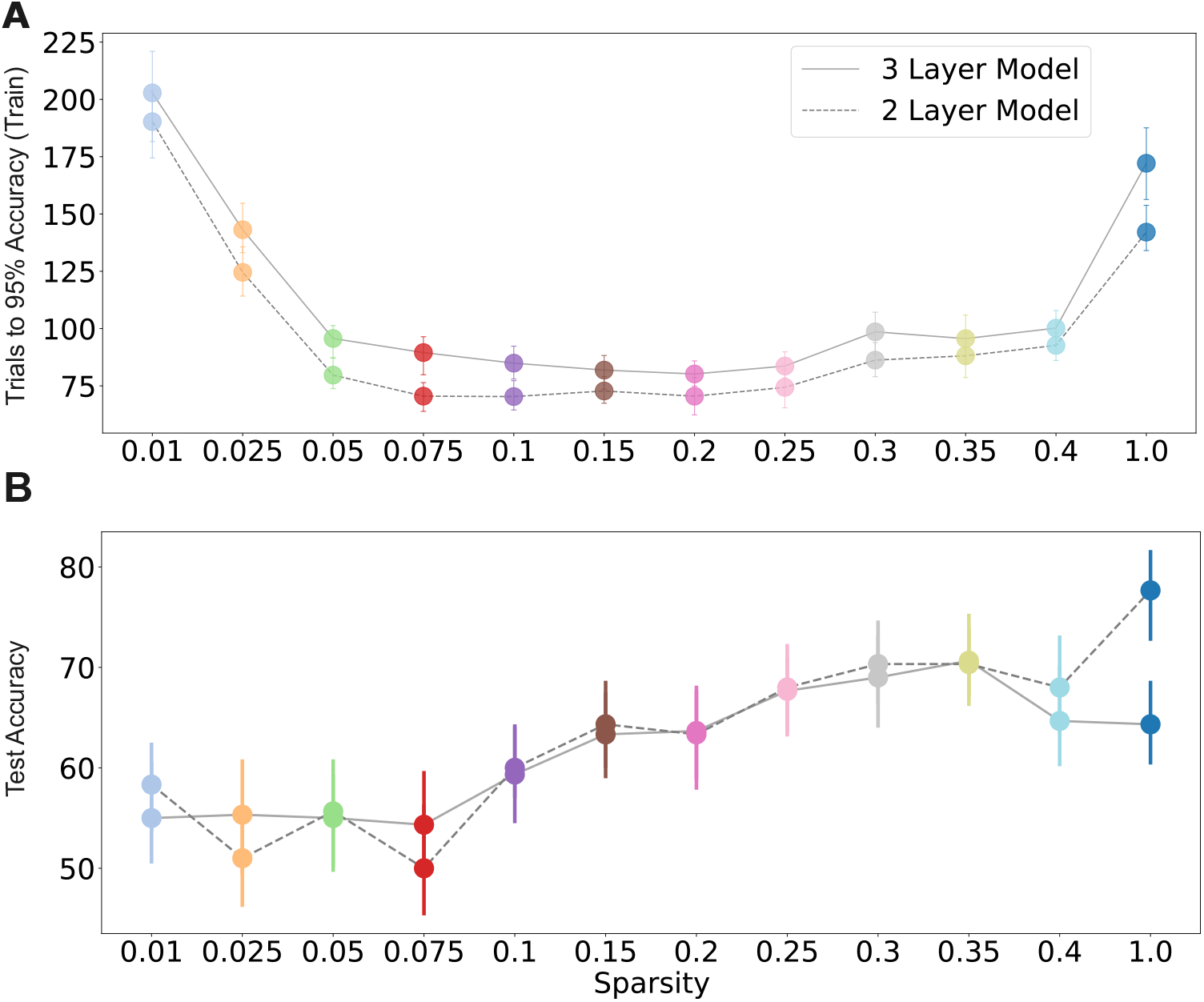
Sparsity level differentially affects learning speed and generalization. (A) Mean number of training trials required to reach 95% accuracy as a function of sparsity level for complete sparse two- and three-layer models trained on the 7-D binary classification task. Models exhibit a non-monotonic dependence on sparsity, similar to the XOR task. (B) Test set accuracy as a function of sparsity level. Generalization improves with denser codes, in contrast to the non-monotonic dependence of training speed on sparsity.

Next, we trained models with sparsity implemented at different positions within the network, with sparsity fixed at 5%. The pattern of learning dynamics across models on the training set closely mirrored our findings in the XOR task (Fig.4A-B, left). Models that had sparsity implemented in the first layer learned fastest. These were followed by models with sparsity confined to the second and third layers, and finally by dense models. Test accuracy revealed a remarkable difference between input-sparse and later-layer-sparse models (Fig.4A-B, right). Models with first-layer sparsity generalized the worst, achieving accuracies around chance level. Models with sparsity implemented in second and third layers exhibited the fastest and strongest generalization performance, achieving test accuracies of ∼ 80-95% (Fig.4(C-D)). The dense model generalized less effectively, reaching ∼ 65-75%. This indicates that first-layer sparsity promotes fast learning through rapid memorization, at the cost of poor generalization.

**Figure 4:**
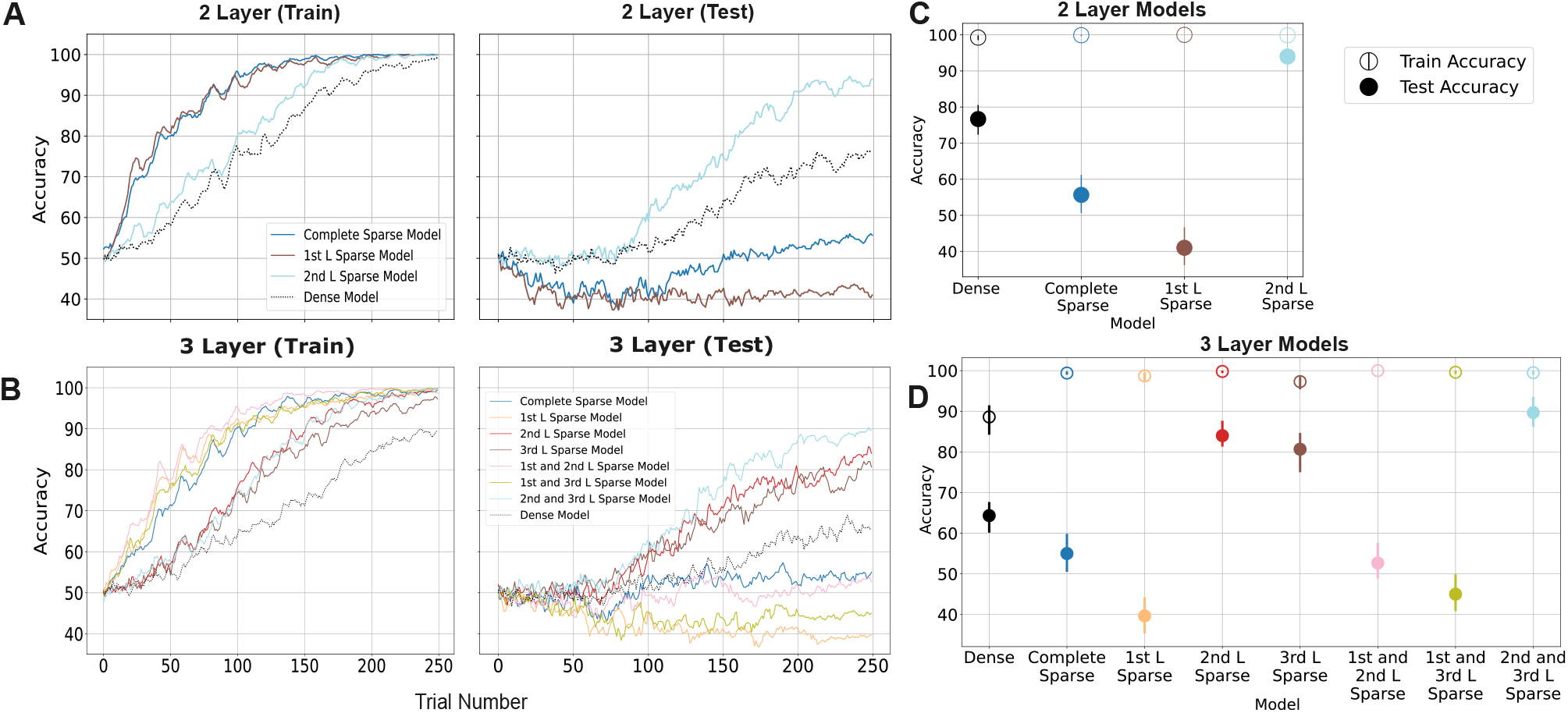
Sparsity at the input versus later stages determines whether models memorize or generalize. (A-B) Learning curves on the training set (left) and test set (right) for two-(A) and three-layer (B) models trained on a 7-dimensional category learning task. Models with first-layer sparsity exhibit fast learning (training set) but poor generalization performance. Models with later-layer sparsity exhibit the best generalization performance (test set). (C-D) Final post-training accuracy (train and test) for two-(C) and three-layer (D) models. Although all networks achieve near 100% training accuracy, test accuracy diverges across models. Models with sparsity confined to later layers generalize best, followed by the dense baseline, whereas model with first-layer sparsity exhibit rapid memorization but poor generalization. Error bars represent bootstrapped 95% CIs.

We performed dimensionality and representational similarity analysis as above and found consistent results to those in the XOR task, complemented by additional insights from the test set (Fig.S2). In the first layer, first-layer sparse and complete sparse models exhibited higher dimensionality than second-layer sparse and dense models, as in the XOR task, except that overall dimensionality values were higher. In the second layer, all models began with relatively high PR values and slowly declined over learning. The higher overall dimensionality likely reflects the more complex task structure. Nevertheless, the pattern is consistent with the XOR task. The first-layer sparse and complete sparse models exhibited the highest dimensionality overall, and the second layer of the first-layer sparse model exhibited high dimensionality which then dropped rapidly, reflecting the adaptation of its internal representations to a low-dimensional space focused on encoding the examples presented in learning. Across the 2D and 7D tasks, sparse architectures exhibited higher initial PR values in layers where sparsity was implemented (Fig.2C; Fig.S2), consistent with an orthogonal, high-dimensional coding regime.

Representational similarity analyses on the training set showed greater within-class than between-class similarity in the second and output layers, consistent with the XOR task. First-layer sparse and complete sparse models exhibited earlier within- and between-class separation in second and output layers compared to the second-layer sparse and dense models. Similar to the XOR task, the dense model began with higher overall similarity values, reflecting more overlapping representations, which gradually separated over learning.

Similarity results on the test set revealed interesting differences across models (Fig.S3). While first-layer sparse and complete sparse models exhibited strong separation on the training set, only the second-layer sparse model showed robust divergence between within- and between-class similarity at the output layer on the test data, closely tracking successful generalization performance. The dense model showed a weaker and later separation, and first-layer and complete sparse models showed none, reflecting memorization of the training set. In sum, representational similarity analyses showed that first-layer sparse models memorize quickly but cannot generalize, whereas second-layer sparse models learn more slowly and form separated representations of new examples.

## 4 Discussion

In this work, we provide a systematic demonstration of how the computational properties of the DG-like sparse code enable fast learning on complex tasks relevant to higher cognition. We show that learning speed varies non-monotonically with sparsity, peaking at a capacity-sufficient level, and that sparsity at the input stage, as in the DG, is required for fast learning.

However, input-stage sparse models learn fast by memorization — through high-dimensional, pattern-separated representations — suggesting the DG sparse code supports rapid encoding of specific experiences, rather than the extraction of generalized knowledge.

Prior work has largely focused on memory function([25]) and simple linear tasks([27]). Models incorporating the DG have shown that a sparse code orthogonalizes patterns([14]), but have not systematically tested how it affects learning. That DG sparse coding facilitates fast learning has therefore been assumed rather than demonstrated. Here, we characterize the conditions under which sparsity helps or hurts learning speed, and show that the sparse DG code produces fast learning on complex, non-linear tasks requiring the flexible, structured learning the hippocampus is thought to support([21]). An extreme sparse code restricts representational capacity, whereas a dense code learns slowly, and the models that learn fastest have sparsity levels similar to the empirical estimates for the DG.

Why does sparsity speed learning? The extreme sparse models have too few active units to update weights as flexibly as larger models — much like training a very small network. A single hidden layer with two units is sufficient to solve the XOR task, but this network has only nine trainable parameters and is hard to train, as both units must converge on complementary solutions. Additional units enlarge the set of solutions, making one easier to reach from a random initialization and allowing it to be distributed across the layer. Extreme sparsity creates a similar bottleneck: although the layer has 200 units, only a handful are active and updated on each trial. At the other extreme, every input in a dense network drives the entire layer, so activations and weight updates from different inputs overlap and interfere. Learning is slow because each step must resolve updates that pull shared weights in conflicting directions. A sparse network with sufficient capacity activates a fixed number of units on each trial, and therefore only updates a small proportion of the weights. Early in learning, as the network is still configuring, a given input may recruit different sets of top-k units, since small differences in weights determine which units win the k-WTA competition. As learning stabilizes, the same units are consistently active for a given input, and only their associated weights are trained. Across the sparsity range, learning speed increases sharply once the layer holds enough units to learn efficiently, then declines gradually as codes become denser and interference builds (Fig.1C). The optimum sits near the sparse end of the range, at sparsity levels matching the DG. Because we mapped learning across the full range of sparsity levels rather than fitting sparsity to data([39]), we characterize how learning speed changes across the range, not only where it peaks.

We found that all first-layer sparse models learn fast, whereas all second- and third-layer sparse models learn markedly slower, demonstrating that sparsity implemented at the input stage of the processing stream is key. This is exactly where the DG sits in the hippocampal trisynaptic circuit, suggesting its position at the circuit’s entry is necessary rather than incidental to fast learning. Notably, the first-layer sparse and first-and-second-layer sparse models match sparse codes found in DG and CA3([44]). Pattern separation is attributed primarily to the DG and its mossy fiber projection to CA3, rather than CA3 itself, matching our finding that models with sparsity in the second layer by itself do not show the same speedup in learning. Why does input-stage sparsity speed learning? Simply having fewer weights to train is not sufficient, or else we would expect models with later sparse layers to learn as quickly. Instead, a sparse first layer performs pattern separation, reducing overlap across inputs and leaving the following layers a simpler problem to solve. A dense first layer exhibits overlapping representations that take many trials of learning to resolve, and later layers inherit this overlap.

Second- and third-layer sparse models learn faster than fully dense models, consistent with fewer trainable weights giving a partial benefit. Notably, they learn at a similar speed to each other and slower than first-layer sparse models, again demonstrating that sparsity at the input stage is what drives fast learning. Sparsity at the input stage speeds learning through pattern separation, but it is precisely this removal of overlap that discards the shared structure across inputs that enables generalization.

Our dimensionality analysis supports this account. In two-layer models, first-layer sparse and complete sparse models show markedly higher dimensionality in the first layer than second-layer sparse and dense models, reflecting more orthogonalized, pattern-separated representations. This is present from the start and across learning, indicating that the separation is imposed by the sparsity mechanism regardless of the task. In the second layer, dimensionality in first-layer sparse models falls sharply early in learning, as the dense layer settles into a low-dimensional readout of an already-separated code. By contrast, in dense and second-layer sparse models dimensionality rises over learning. Complete sparse models show no such drop, as sparsity is implemented in the second layer.

We find that sparsity trades generalization for speed, with input-stage sparse models learning fast through memorization. Where sparsity is implemented determines which side of the memorization–generalization spectrum a model falls on, while the level of sparsity modulates the effect. Generalization improves as codes become denser, in contrast to the inverted-U for learning speed.

At matched sparsity, models with sparsity implemented in the first layer memorize while those with sparsity in later layers generalize, showing that input-stage sparsity restricts generalization independently of sparsity level. Examining the internal representations revealed that only later-layer sparse models develop separated category representations for test examples, whereas early-sparse models separate the training examples, corresponding to generalization performance.

Storing specific experiences is a large part of what the hippocampus does, and a code that keeps similar experiences distinct is well suited to that pur ose. Pattern separation is what makes episodic storage possible, and what limits generalization. The two objectives need not be served by the same code: the hippocampal sparse code retains specific experiences and the cortical dense code extracts regularities across experiences([45]). Our contribution is that a code’s position on the memorization–generalization spectrum is set by two properties: how sparse it is, and where sparsity is implemented. Only models with input-stage sparsity produce fast, memorization-based learning – a constraint not previously described. Notably, the hippocampus itself supports generalization and inference([46]; [47]), which under our account arise not from the DG’s representations but from mechanisms elsewhere in the circuit.

Work on overparameterization and double descent has shown that where a network sits on the memorization– generalization spectrum depends on capacity, not on the data or objective alone([35]; [36]; [37]). Sparsity shifts a network along the same spectrum by a distinct route, constraining how many units are active rather than how many the network has. It also introduces a dependence on position that this literature does not address: the same constraint at different stages produces opposite outcomes. Specifically, at fixed capacity, position determines whether a network memorizes or generalizes, and level determines the degree to which it does so.

Our account leaves open how the hippocampus achieves generalization. The circuit contains pathways that bypass the DG, from entorhinal cortex to CA3 and to CA1([48]). It is also highly recurrent, through CA3’s recurrent collaterals and big-loop recurrence returning hippocampal output to entorhinal cortex([49]). Prior work has shown that recurrence supports inference across related episodes([50]; [51]), though this concerns inference over memories rather than generalization from learned representations. C-HORSE, a hippocampal model, can learn categories and generalize to novel exemplars([52]). In that model, the sparse trisynaptic route encodes individual items and the denser monosynaptic route extracts category structure, though what determines this functional split remains open. Our results suggest one part of the answer: where sparsity sits in the processing stream determines whether a route memorizes or generalizes. Systematically varying these pathways and recurrent connections would establish which support hippocampal generalization.

The DG’s sparse code and its position at the entry to the trisynaptic circuit underlie our capacity to learn rapidly from individual experiences. These same properties make the DG’s code a mechanism suited to forming episodic memories rather than extracting general structure. We provide a principled account of when hippocampal sparse coding helps learning and when it hurts it, showing why sparsity at the circuit’s first stage commits the DG to separated representations that serve not only memory but the wider range of complex cognition the hippocampus supports.

## 5 Methods

### 5.1 Experiment and simulation description

We investigated how varying levels of hippocampus-like sparsity affect learning speed and generalization. To this end, we constructed a family of models differing only in the degree of enforced sparsity and compare their performance against a non-sparse (dense) baseline.

### 5.2 Modeling hippocampal sparsity

To model the sparse activity patterns observed in the hippocampus, we implemented a *k*-winners-take-all (*k*-WTA) mechanism([53]0; [25]), which approximates the effect of inhibition by hippocampal interneurons([54]). For a population of *N* units with activations *a* ∈ ℝ^*N*^, activity was restricted to the top *k* units for a given input, with all remaining units set to zero. Sparsity was defined as the proportion *p* of units active for a given input, so that *k* = *pN*. Both were fixed within a given model, except in the width control, where *k* was instead held constant across layer widths (see Experiments).

We used standard multilayer feedforward neural networks as a controlled and interpretable framework for studying the effects of sparsity. These models provide a setting in which architectural and functional properties can be systematically manipulated, allowing precise control over sparsity through *k*-winners-take-all inhibition. This enabled us to vary sparsity independently of other factors and isolate its impact on learning dynamics and generalization.

This served as a simplified model of hippocampal processing to focus on the computational role of sparsity itself. This allowed us to directly examine how sparse activity patterns, akin to those observed in hippocampal regions such as the dentate gyrus, influence representation formation and learning efficiency. As such, the multilayer network provided a tractable and principled testbed for studying sparsity as a computational mechanism, while abstracting away secondary biological effects.

Let *x* ∈ ℝ^*d*^ be the input to a layer, *W* ∈ ℝ^*N* ×*d*^ and *b* ∈ ℝ^*N*^ the learnable parameters, and *ϕ*(·) an element-wise activation function. The layer computed

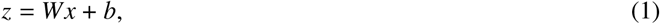

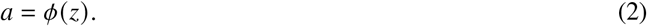

Let *S*_*k*_ (*a*) denote the index set of the *k* largest components of *a*. Define a binary mask **m** ∈ {0, 1}^*N*^ as

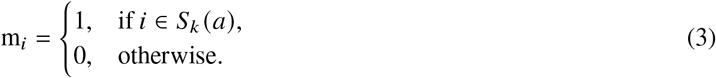

The output of the layer was

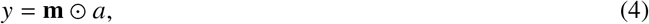

where ⊙ denotes element-wise multiplication. If *k* = *N*, the operator reduces to the identity and *y* = *a*. Equivalently, the layer can be written compactly as

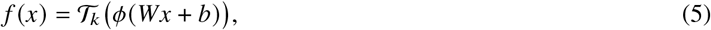

where *T* _*k*_ (·) denotes the top-*k* selection operator that keeps the *k* largest components and sets the rest to zero. A dense layer omits this operation, yielding

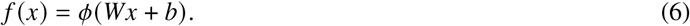

### 5.3 Random-*k* sparsity control

To distinguish the effects of structured sparsity from those of reduced activity alone, we additionally implemented a *random-k sparsity* control condition. In this setting, instead of selecting the top-*k* units based on activation magnitude, a random subset of *k* units was selected independently for each forward pass, with all remaining units set to zero.

Formally, let *R*_*k*_ ⊂{1, …, *N*} denote a randomly sampled subset of size *k*. The binary mask **m** ∈{0, 1}^*N*^ is then defined as

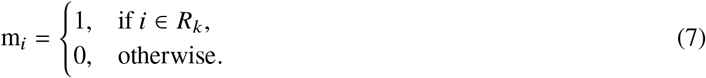

The layer output was *y* = **m** ⊙ *a*, analogous to the *k*-WTA case but without dependence on the activation values. This control preserved the same sparsity level while removing competitive selection, isolating the role of structured, activity-dependent sparsification.

### 5.4 Training and model evaluation

Models were trained using a binary cross-entropy loss, appropriate for the binary classification setting considered here. Given a model output probability *p* ∈ (0, 1) and binary target label *y* ∈ {0, 1}, the loss is defined as:

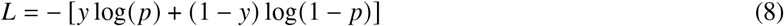

Optimization was performed using the Adam optimizer. Learning rates of 0.01 for the 2D input-size problem and 0.005 for the 7D input-size problem were selected to ensure stable learning dynamics and reliable convergence to asymptotic performance (near 100% accuracy) without training instability or premature stagnation. Models were trained with a batch size of 1, such that parameters were updated after each trial, consistent with human learning.

To evaluate model performance, we used the percentage accuracy score. The accuracy score computes the number of correct classifications as a ratio of the total trials reported as a percentage.

Model performance was evaluated using classification accuracy, computed as the percentage of correctly classified trials. Models were trained over 50 independent runs. Reported accuracies per trial correspond to the mean percentage correct across all inputs (the full training set or the full test set, respectively) across runs, and 95% confidence intervals were estimated using a non-parametric bootstrap procedure with resampling over runs (100 bootstrap samples).

### 5.5 Representational similarity analysis

We performed a form of representational similarity analysis (RSA)[55] to evaluate the models’ representations of task-relevant information by averaging cosine similarity within and between classes. For a given layer, unit activations were extracted for each trial, and pairwise cosine similarity was computed between activation vectors. Within-class similarity was computed as the mean cosine similarity over all pairs of inputs belonging to the same class. Between-class similarity was computed as the mean cosine similarity over all pairs of inputs across different classes.

Let 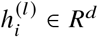 denote the representation of trial *i* at layer *l*, and let *C* index classes, with *I*_*c*_ the set of trials belonging to class *c*. Define cosine similarity as

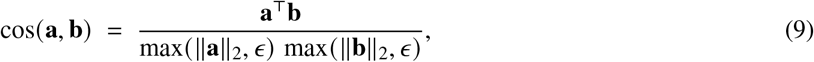

where *ϵ* = 10^−8^. Within-class representational similarity at layer *l* was then the mean cosine similarity over all pairs of trials belonging to the same class,

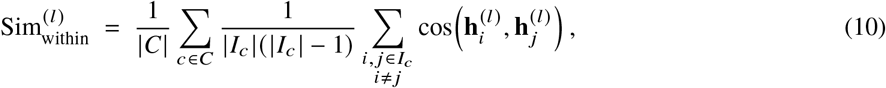

and between-class similarity was the mean cosine similarity over all pairs of trials belonging to different classes,

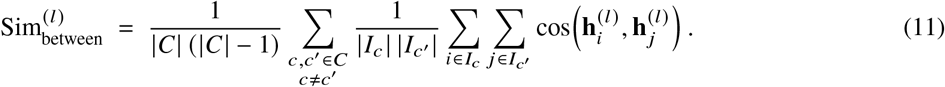

Both measures were computed at each layer and at each point in training.

### 5.6 Dimensionality analysis

To quantify the dimensionality of representations across layers, we computed the participation ratio (PR) of unit activations[43]. PR provides a measure of how many dimensions are used to encode information.

Let 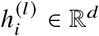 denote the activation vector (i.e., representation) of the *i*-th input at layer *l*, and let *X* ^(*l*)^ ∈ R^*n*×*d*^ be the matrix of activations across *n* trials. We first centered the representations by subtracting the mean across trials:

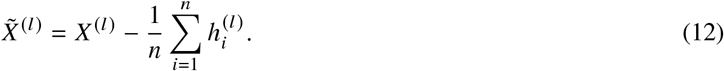

We then computed the covariance matrix

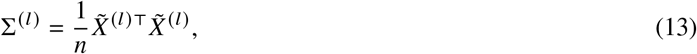

and let 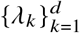 denote its eigenvalues. The participation ratio at layer *l* is defined as

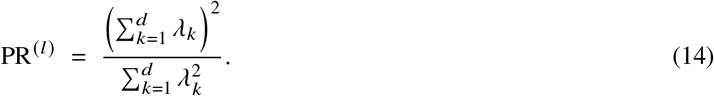

The participation ratio can be interpreted as the number of dimensions used by the representation. If variance is evenly distributed across many directions, PR is high, indicating a high-dimensional code. Conversely, if variance is concentrated along a few directions, PR is low, reflecting a low-dimensional representation. We used PR to track how sparsity shapes the dimensionality of representations during learning.

### 5.7 Model setup

The baseline architecture was a fully-connected feedforward network with either two or three hidden layers followed by an output layer. Hidden layers contained 200 units each, and the output layer consisted of 2 units corresponding to the binary classification task. All models shared identical weight initialization, activation functions, and training procedures, differing only in the application of sparsity.

Hidden-layer activations used a LeakyReLU nonlinearity with a negative slope set to 0.1. This choice was motivated both by its closer correspondence to biological neurons, which exhibit non-zero baseline activity, and by its empirical stability under trial-by-trial learning. In particular, compared to ReLU, tanh, sigmoid, and softplus activations, LeakyReLU consistently yielded faster and more reliable convergence when models were trained with one example per trial. Output-layer activations are as described in the modeling sparsity section.

Sparsity was implemented at the level of hidden-layer activations using the *k*-winners-take-all mechanism described in the previous section. When a layer was designated as sparse, only the top 5% of units (corresponding to *k* = 10 active neurons out of 200) were permitted to remain active for a given input, with all other activations set to zero. Dense layers retained all activations without additional inhibition.

To assess the effect of sparsity placement, we evaluated a set of model variants corresponding to different configurations of sparse and dense hidden layers:

1. a fully dense model (no sparse layers),
2. a model with sparsity applied only to the first hidden layer,
3. a model with sparsity applied only to the second hidden layer,
4. a model with sparsity applied to both hidden layers.

For three-layer models, all eight combinations of sparse and dense hidden layers were evaluated.

All models contained the same total number of parameters prior to sparsification, ensuring that performance differences arose from the presence and position of sparsity constraints.

In addition, to examine whether the effects of sparsity placement persisted with increased network depth, we performed supplementary experiments on models with a larger number of hidden layers. The results of these experiments are reported in the Supplementary Material.

All models were implemented in PyTorch (v2.6.0). Numerical analyses were performed using NumPy (v2.2.6), SciPy (v1.16.3), pandas (v2.3.3), and scikit-learn (v1.8.0). Dimensionality was quantified using the participation ratio as implemented in scikit-dimension (v1.8.0). Figures were generated using Matplotlib (v3.10.6) and Seaborn (v0.13.2).

### 5.8 Experiments

We evaluated the impact of sparsity on learning speed and generalization across several controlled experimental settings.

We first investigated how different sparsity levels affect learning dynamics. We varied the sparsity proportion *p* ∈{0.01, 0.025, 0.05, 0.075, 0.1, 0.15, 0.2, 0.25, 0.3}, where sparsity was implemented by allowing only the top-*p* × *N* neurons (based on activation magnitude) to remain active in a layer, with all others suppressed. Here, *N* = 200 was the default hidden-layer width used in both two- and three-layer models.

To examine how network width interacts with sparsity, we trained models with varying hidden-layer sizes *N* ∈ {100, 200, 300, 400, 1000} under two sparsity regimes: (i) *percentage-based sparsity*, where the top 5% of units were active, and (ii) *absolute sparsity*, where a fixed number of units (*k* = 10) were active, corresponding to the active set size for the default width of *N* = 200. This allowed us to disentangle the effects of the absolute number of active units from the proportion of active units.

To study how the position of sparsity within the network influences learning, we trained two- and three-layer architectures with different configurations of sparse and dense layers. In these experiments, hidden-layer width was fixed at *N* = 200, and sparsity was set to *p* = 0.05 (that is, top-10 active units in sparse layers). For two-layer models, we considered four configurations: fully dense (no sparsity), first-layer sparse, second-layer sparse, and fully sparse. For three-layer models, we evaluated all permutations of sparse and dense layers.

All experiments described above were conducted on a two-dimensional XOR classification task to study learning speed in a controlled setting. To evaluate generalization, we further considered a higher-dimensional rule-plus-exception task with 7-dimensional inputs and a train–test split. In this setting, we assessed the performance of different sparsity configurations in both two- and three-layer models on held-out data. For these experiments which involve a train–test split (7-D rule-plus-exception task), generalization performance was evaluated throughout training by periodically measuring accuracy on the held-out test set. Importantly, test-set evaluation was performed in a purely feedforward manner without updating model weights, ensuring that test performance reflected true generalization rather than additional learning.

## 6 Acknowledgments

We thank CiNet members for constructive feedback on earlier presentations of this work.

## 7 Data availability

The datasets generated and analyzed in this study are available at [OSF Link](https://osf.io/a42t9/overview?view_only=8fa7b84c68214235a9aa69f5c571084c)

## 8 Code availability

All code used to generate the results in this study is available at [Github Link](https://github.com/animeshsasan/hippocampal-cortex-learning).

## 9 Supplementary information

### 9.1 7D category learning task

**Figure S1:**
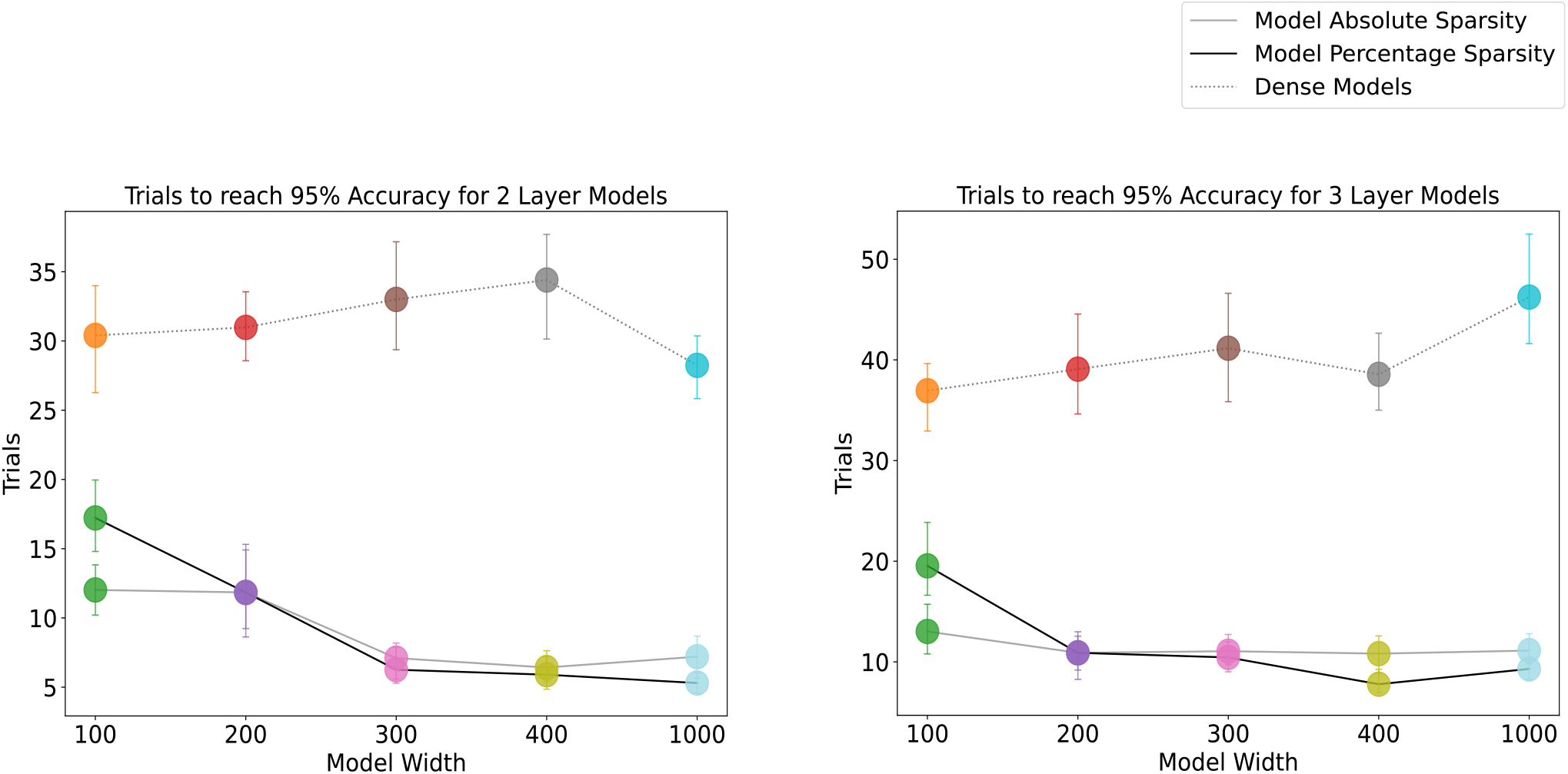
Effect of network width on learning speed under different sparsity regimes. Number of training trials required to reach 95% accuracy as a function of hidden-layer width for (A) two-layer and (B) three-layer architectures. Results are shown for dense models (dotted line), percentage-based sparsity (black line), and absolute sparsity (gray line).

**Figure S2:**
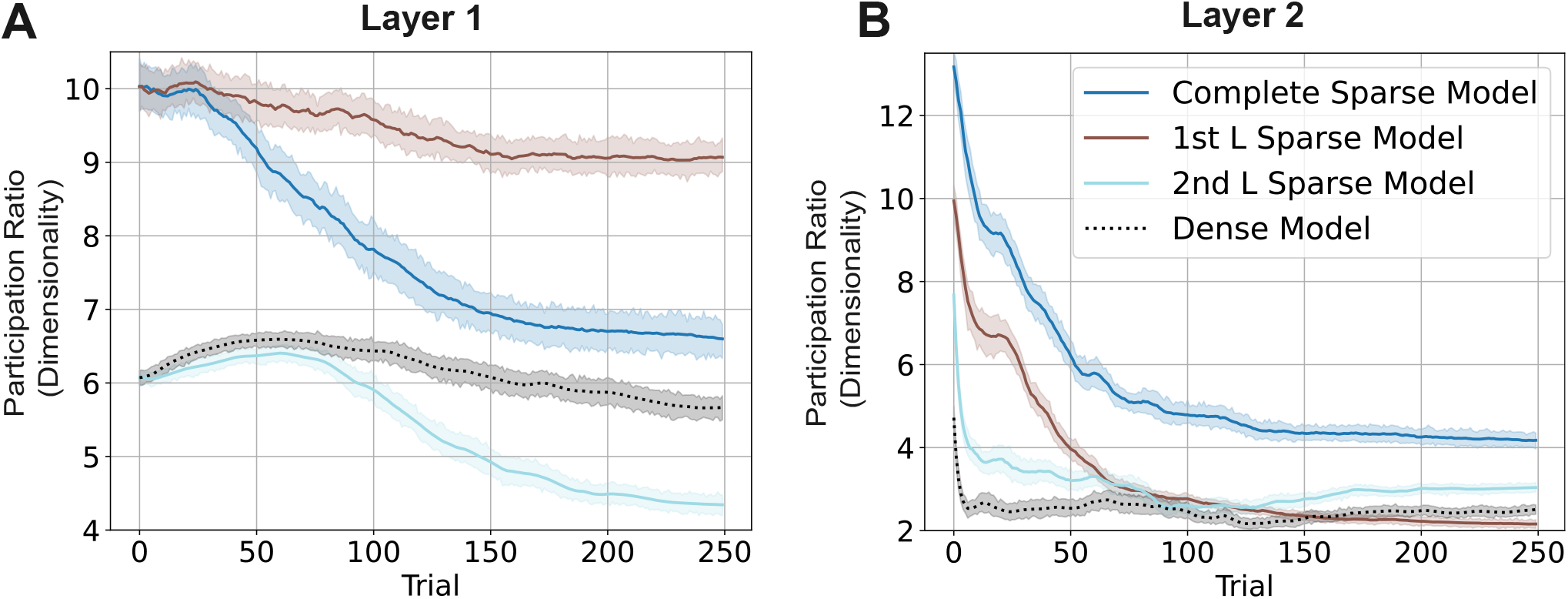
Dimensionality dynamics across layers in the 7-D task. Participation ratio (PR) is shown across training trials for two-layer models trained on the 7-D binary classification task. Results are plotted for the first hidden layer (A), and second hidden layer (B) for Dense, Complete Sparse, 1st-Layer-Sparse, and 2nd-Layer-Sparse architectures. Shaded error bars represent 95% bootstrapped CIs.

**Figure S3:**
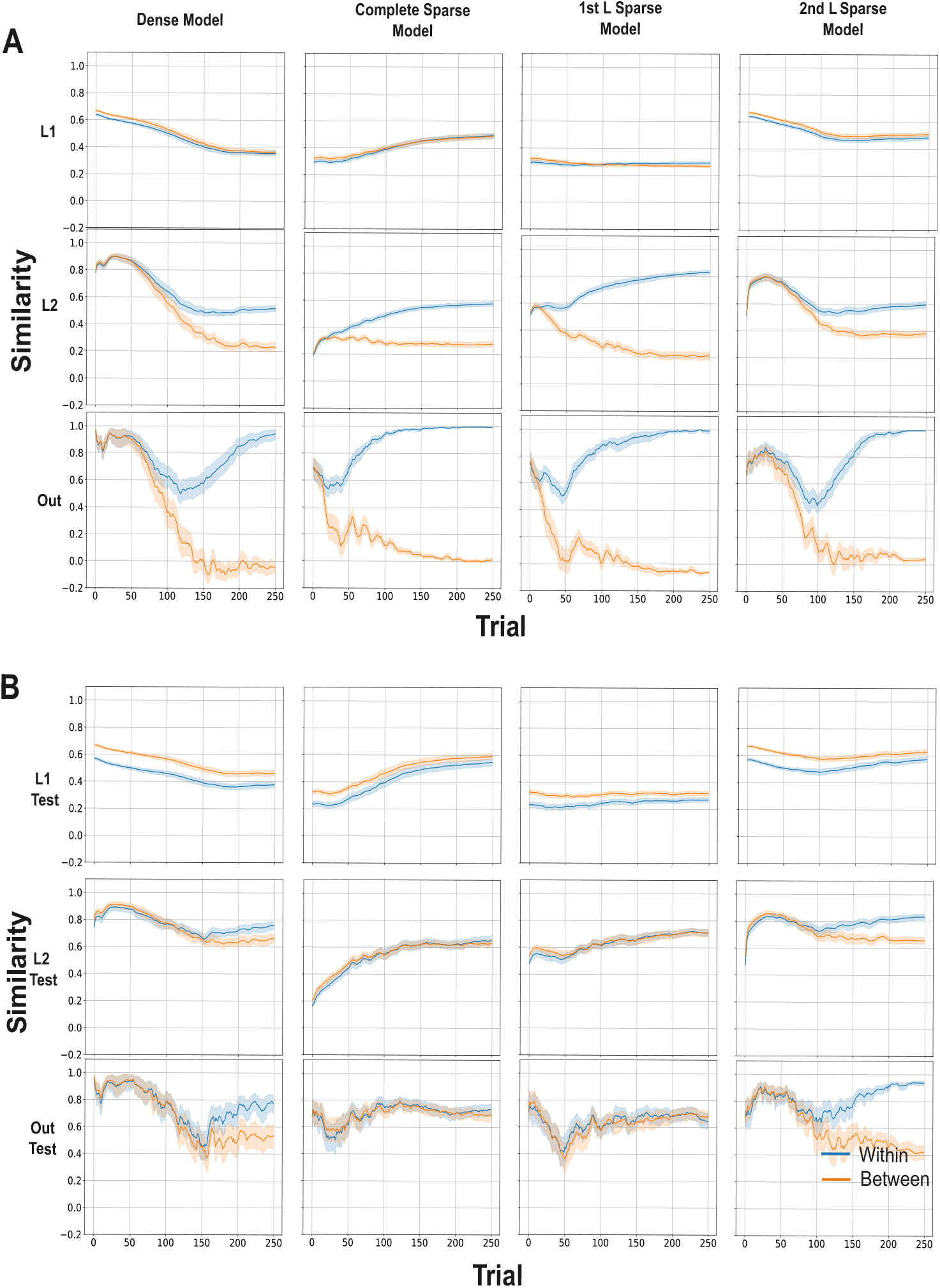
Layer-wise representational similarity dynamics for 2-layer models on the 7-D task. Cosine similarity between activations, averaged across within-class (blue) and between-class (orange) input pairs, is shown across training trials for the train set (A) and test set (B). Results are plotted separately for layer-1 (L1), layer-2 (L2), and output (Out) representations in dense, complete sparse, first-layer sparse, and second-layer sparse models.

**Figure S4:**
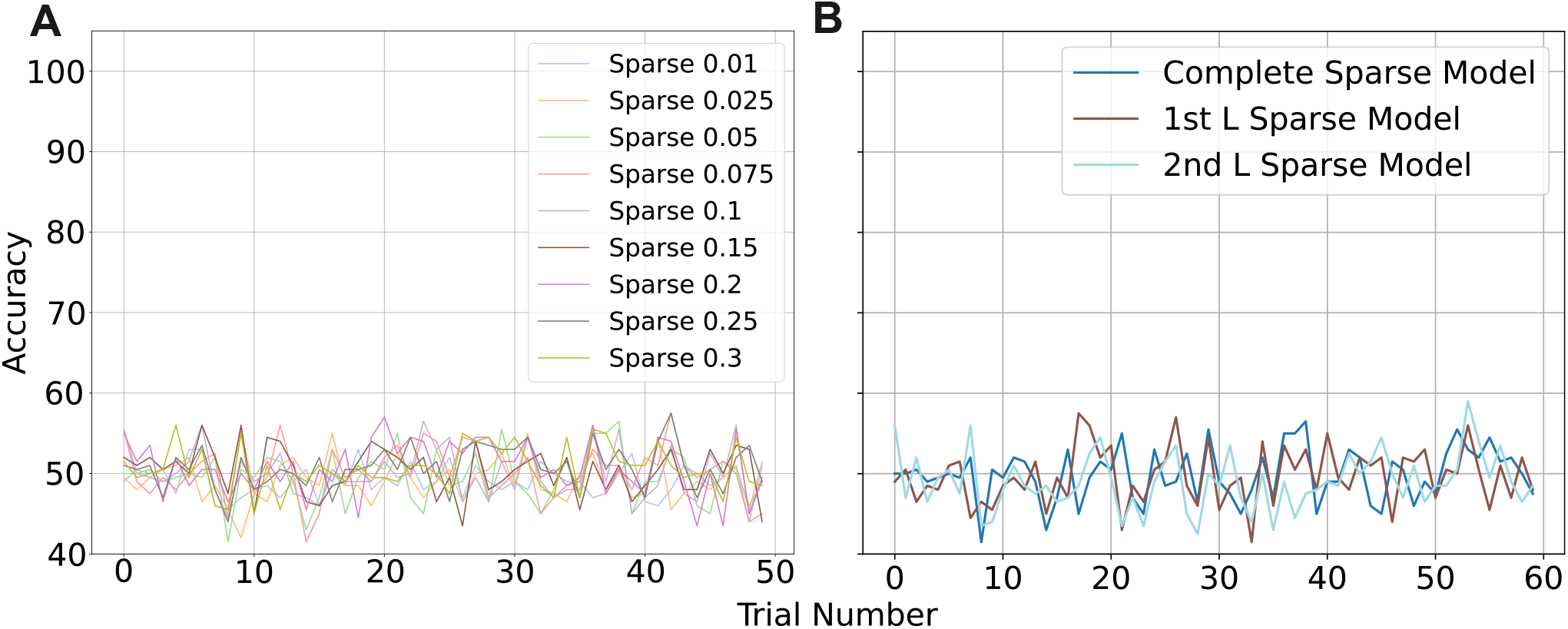
Effect of random-k sparsity on learning dynamics in two-layer networks. Learning curves for two-layer networks trained on a 2D XOR task implemented with random-*k* sparsity levels (A) and sparsity (5%) implemented at different layers (B).

